# Low Computational-cost Cell Detection Method for Calcium Imaging Data

**DOI:** 10.1101/502153

**Authors:** Tsubasa Ito, Keisuke Ota, Kanako Ueno, Yasuhiro Oisi, Chie Matsubara, Kenta Kobayashi, Masamichi Ohkura, Junichi Nakai, Masanori Murayama, Toru Aonishi

## Abstract

The rapid progress of calcium imaging has reached a point where the activity of tens of thousands of cells can be recorded simultaneously. However, the huge amount of data in such records makes it difficult to carry out cell detection manually. Consequently, because the cell detection is the first step of multicellular data analysis, there is a pressing need for automatic cell detection methods for large-scale image data. Automatic cell detection algorithms have been pioneered by a handful of research groups. Such algorithms, however, assume a conventional field of view (FOV) (i.e. 512 × 512 pixels) and need a significantly higher computational power for a wider FOV to work within a practical period of time. To overcome this issue, we propose a method called low computational-cost cell detection (LCCD), which can complete its processing even on the latest ultra-large FOV data within a practical period of time. We compared it with two previously proposed methods, constrained non-negative matrix factorization (CNMF) and Suite2P. We found that LCCD makes it possible to detect cells from a huge-amount of high-density imaging data within a shorter period of time and with an accuracy comparable to or better than those of CNMF and Suite2P.

## 1. Introduction

The activities of many nerve cells can now be measured simultaneously for a long time, and with this capability, long-term temporal changes in the information representation of a neuronal population and the functional interactions within it can be investigated (Ziv et al., 2013). The rapid progress of imaging technology has made it possible to carry out such research. For example, various high-efficiency fluorescent calcium indicator proteins have been developed, including GCaMP with high calcium sensitivity (Chen et al., 2013). Regarding progress in imaging devices, a miniaturized microscope mounted on the head of a mouse has enabled long-term measurements of neural population activities during long-term behaviors and learning (Ziv et al., 2013). Furthermore, several research groups have developed very wide field-of-view two-photon microscopies capable of imaging areas of square millimeter order (Stirman et al., 2016; Sofroniew et al., 2016). Such ultra-large field-of-view microscopies have made it possible to image the activities of several thousand to several tens of thousands of cells at single-cell resolution.

The first step of an analysis of multi-cellular image data is identification of the positions of the individual cells as regions of interest (ROIs) within the image; this step is called ‘cell detection’ (Lutcke and Helmchen, 2011). Manual identification of ROIs is effective, but it needs a lot of time and effort. As the data size becomes larger, manual cell detection is becoming more and more difficult, and the demand for automated cell detection is increasing. Here, several automatic cell detection methods using machine learning have come to be used. For instance, Mukamel et al. (2009) proposed a cell detection method using the principal component analysis (PCA)-independent component analysis (ICA) algorithm. Moreover, several methods using non-negative matrix factorization (NMF) have been proposed. In particular, Maruyama et al. (2014) proposed background-constrained NMF (BC-NMF), which contains a background fluorescence model; it was shown to outperform the PCA-ICA method in noisy situations. Pnevmatikakis et al. (2016) proposed a constrained NMF (CNMF) with an autoregressive model for describing transient changes in neural calcium profiles. CNMF was shown to outperform both PCA-ICA and BC-NMF. Marius et al. (2016) proposed a NMF based method called Suite2P, which contains a model of background fluorescence changes due to neuropil activities. Different from these NMF methods, Reynolds et al. (2017) proposed ABEL, a method based on an active contour model. It has been reported that ABEL outperforms CNMF and is comparable in performance to Suite2P.

Since all the above methods are formulated as large-scale optimization problems, they are very costly in terms of their computation and memory requirements when dealing with large-scale image data. In addition, the number of cells able to be measured at single-cell resolution is rapidly increasing thanks to rapid progress in imaging technology. As demonstrated in this paper, it is difficult for such methods to process large-scale image data acquired with the latest ultra-large field-of-view microscopies within a practical period of time. For this reason, we developed a low computational-cost cell detection (LCCD) method adaptable to large-scale image data acquired with the latest ultra-large field-of-view microscopies. LCCD is a very simple method taking the following temporal division and integration approach: 1) divide up the data into several short time frames; 2) use a simple filter method to detect cells as ROIs from each of the divided data; 3) integrate the ROIs of temporally different data. We confirmed with real data and artificial data that LCCD performed comparably to CNMF and Suite2P, whereas its calculation time was shorter than those of the CNMF and Suite2P. In particular, LCCD could complete a cell detection task using artificial data mimicking data on the scale of ultra-large field-of-view microscopies in a reasonable amount of time, whereas CNMF and Suite2P could not do so.

## 2. Materials and Methods

### 2.1. LCCD

#### 2.1.1. Outline of LCCD

A schematic diagram of LCCD is shown in Fig. 1. LCCD is a very simple method taking the temporal division and integration approach. The main routine is presented in Algorithm 1, and the outline of its operation is as follows; 1) divide up the data into several short time frames; 2) use a blob detector (BD) to pick up cells (as the ROIs) from each of the divided frames; 3) integrate these ROIs by using the index of spatial overlap between temporally different ROIs, and delete ROIs exceeding the size of a single cell.

**Figure 1:**
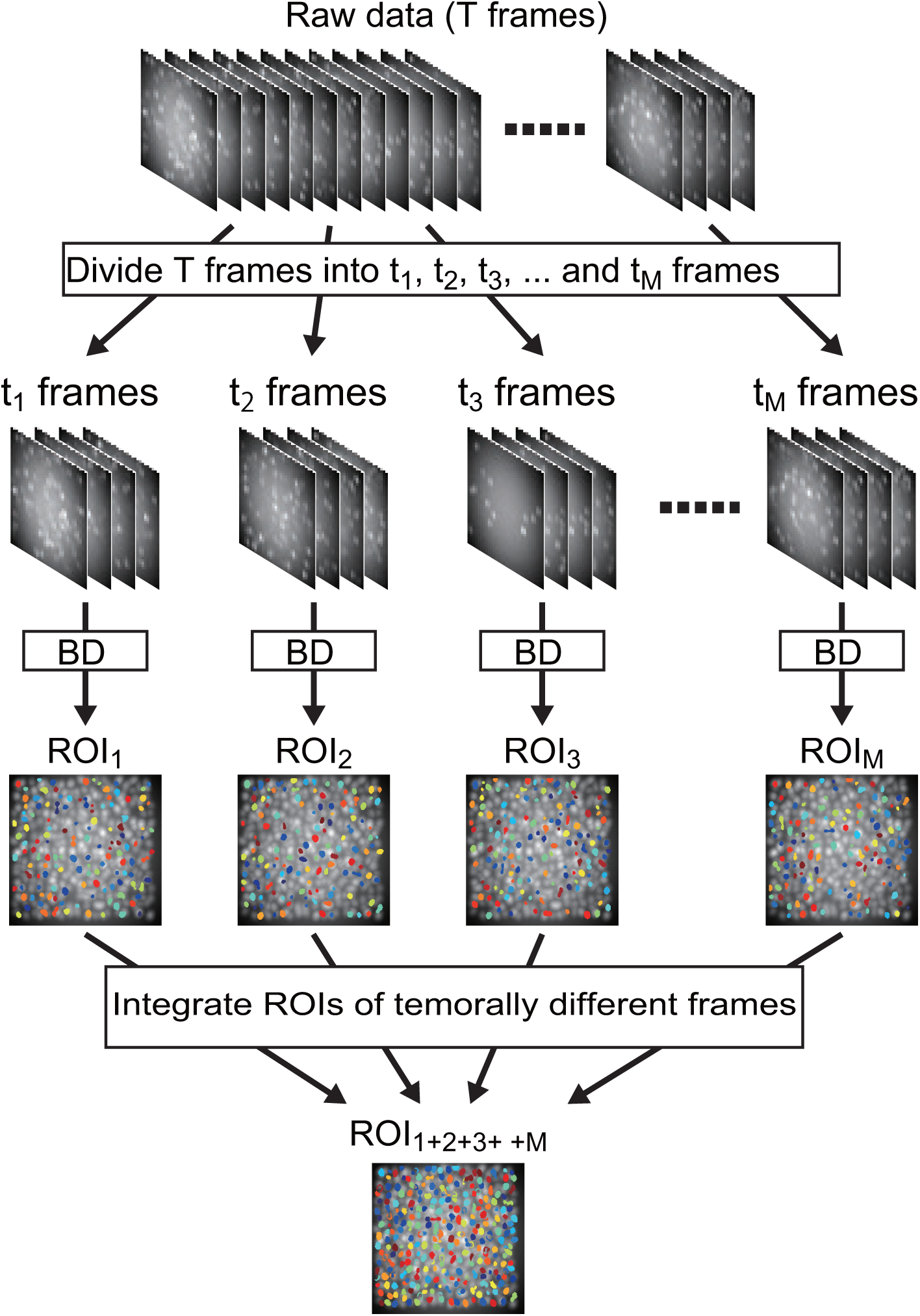
Schematic diagram of LCCD taking temporal division and integration approach: 1) divide up the data into several short time frames; 2) use BD to detect cells as ROIs from each of the divided frames; 3) integrate ROIs of temporally different frames.

#### 2.1.2. Blob detector

The blob detector (BD) is used to detect cells as ROIs from the divided frames. Its routine is presented in Algorithm 2, and its operation is as follows.

(line 2-5 in Algorithm 2) First, to remove slow temporal trends and shot noise from the image data, temporally smoothed data by a long moving average filter with a width of 100 frames are subtracted from a temporally smoothed one by using a short moving average filter with a width of several frames. Then all the negative values of the denoised trend-subtracted data are cut with a half-wave rectifier. The values of the parameters used in the following experiment are given in Algorithm 2.

(line 6-7 in Algorithm 2) Next, the maximum intensity projection along the temporal axis is executed to create a maximum luminance variation map.

(line 8-10 in Algorithm 2) A Mexican hat (MH) filter and positive half-wave rectification are then applied to the maximum luminance variation map to emphasize the elliptical shape of cell bodies and reduce the background luminance. The MH filter is constructed by subtracting a two-dimensional averaging filter from a two-dimensional Gaussian filter. The filter size and standard deviation of the Gaussian are manually adjusted according to the expected cell body size. The values of the parameters used in the following experiment are given in Algorithm 2.

(line 11-12 in Algorithm 2) The contrast of the filtered map is emphasized with contrast-limited adaptive histogram equalization (Zuiderveld, 1994).

(line 13-20 in Algorithm 2) After that, the contrast-emphasized map is binarized with a threshold value determined by the Otsu method (Otsu, 1979), and closed regions are detected with an eight-connected-component labeling operation. Next, only the closed regions having areas ranging from 20 to 50 pixels are picked up, and elongated ROIs are excluded with an oval filter (Algorithm 3).

(oval_filter function in Algorithm 3) The oval filter can be easily realized by measuring the following two geometric characteristics of closed regions with the regionprops function in the MATLAB Image toolbox. One characteristic is the eccentricity of the ellipse that has the same second moments as a closed region. The other is the area ratio between a closed region and an ellipse having the same normalized second central moments as the region. Here, we selected closed regions only satisfying the conditions that the eccentricity is less than 0.99 and the area ratio is less than 1.8. By applying the oval filter to the detected closed regions, somata are identified as oval-shaped ROIs.

#### 2.1.3. ROI integration

ROIs, which are detected from each of the divided frames with the BD, are integrated by using the index of spatial overlap between the ROIs of temporally different frames, and ROIs exceeding the size of a single cell are deleted. The routine of ROI integration is presented in Algorithm 4.

The key points of this algorithm are as follows. If area of overlap between two ROIs is more than 40% of the whole area of either ROI, the larger one is selected and the smaller one is deleted (line 9-14 in Algorithm 4). If area of overlap between two ROIs is less than 40% of the area of each of the ROIs, the overlap region is deleted from both the ROIs (line 15-16 in Algorithm 4). Finally, ROIs smaller than 20 pixels and larger than 500 pixels are deleted (line 21 in Algorithm 4).

### 2.2. Compared cell detection methods

CNMF is an NMF-based method that incorporates an autoregressive model of calcium transient dynamics as a temporal constraint. In the program package, the user can choose one of two methods, greedy or sparse, as preprocessing to roughly detect cells. In this study, we chose greedy, which is recommended to be used for detecting cell bodies in the same situation as this research. The CNMF program package used for comparison was downloaded from https://github.com/epnev/ca_source_extraction (Pnevmatikakis et al., 2016). In the numerical experiments, the parameters of the CNMF (the std of the gaussial kernel, number of components to be found, order of autoregressive system, etc.) were manually set to maximize the performance of its cell detection. For a fair comparison with other methods, we only picked up closed regions having an area of 20 to 300 pixels, and we excluded elongated ROIs with the oval filter used by LCCD.

Suite2P is an NMF method that incorporates a model of background fluorescence changes due to neuropil activities. It has been reported that Suite2P outperforms CNMF. The Suite2P program package used for comparison was downloaded from https://github.com/cortex-lab/Suite2P (Marius et al., 2016). In the numerical experiments, the parameters of Suite2P (ops0.nSVDforROI, ops0.NavgFramesSVD, ops0.diameter, etc.) were manually set so as to maximize its performance.

The numerical experiments did not involve any manual editing of ROIs determined by the various cell detections so that only their abilities would be compared.

### 2.3. Computational environment

All programs of LCCD, CNMF, and Suite2P were implemented in MATLAB (MathWorks). The computer used in the numerical experiments had four 22-core Intel (R) Xeon (R) E-7-8880 V4@ 2.2GHz processors (total of 88 cores/176 threads) with a total of 2TB RAM (HPCTECH). The operating system was a Cent-OS 6.7.

### 2.4. Two-photon calcium imaging data

We performed two-photon calcium imaging of L2/3 cortical neurons (100 - 135 um below the pia) in an awake mouse. The images (1.46 um/pixel) were acquired at 7.5 Hz using a custom-modified multiphoton microscope. The neurons expressed calcium sensors GCaMP6f (Chen et al., 2013), GCaMP7.09 or jGCaMP7f (Shiba et al., 2016; Dana et al., 2018) mediated by adeno-associated virus.

### 2.5. Synthesized artificial data

To quantitatively evaluate the performance of LCCD and other methods, we synthesized imaging data of high-density overlapping cells (256 × 256 pixels, 2000 frames). The number of cells was 324. Each cell with an oval shape was placed on a point slightly randomly shifted from a two-dimensional lattice, and its orientation was randomly determined. Each cell was created using a two-dimensional Gaussian with two standard deviations of 13 and 13 pixels. Calcium transients in each cell were mimicked by an exponential decay function with a time constant of 50 frames, and its initializing time point obeyed a simple Bernoulli process with a probability of initialization equal to 0.25 % per frame. For simplicity, the time constant of the exponential decay functions, the maximum rising amplitudes and frequencies of transients were assumed to be uniform in all cells and all frames. Static background fluorescence mimicked by a two-dimensional Gaussian function was added to these cells’ signals. Here, the maximum rising amplitude of each transient and the maximum background fluorescence were each kept constant at 1000. Lastly, artificial movies were synthesized through a shot noise process obeying a Poisson distribution whose intensity parameter was set to be the signal amplitude of each pixel.

Furthermore, to evaluate the calculation time of the methods, we synthesized artificial movies mimicking large-scale calcium-imaging data. We generated four movies with the following scales: 225 cells in 256 × 256 pixels, 841 cells in 512 × 512 pixels, 3136 cells in 1024 × 1024 pixels and 10000 cells in 2048 × 2048 pixels. Each of the four movies consisted of 5000 frames.

### 2.6. Statistical Measures in the Performance Evaluation

To quantitatively compare the performance of the automatic cell detection methods, we used both synthesized data and in-vivo data. Here, the true ROIs were created artificially and used as the ground truth (GT) in the evaluations with the synthesized data, whereas the ROIs obtained with CNMF were used as the GT in the evaluation with in-vivo data. We quantified the level of matching between the GT ROIs and the automatically obtained ROIs by using the following statistical measures.

First, temporal cross-correlation coefficients between fluorescence signals, i.e., df/f of all possible pairs of GT ROIs and ROIs obtained with the other methods, were calculated, and all such pairs were sorted in descending order of the cross-correlation coefficient. Next, the best matched pair of a GT ROI and ROI of another method was selected; then, the next best matched pair, which did not contain ROIs in the previously selected ROI pair, was selected. This selection procedure was repeated until there were no pairs with non-zero correlation.

If the ROI obtained with another method was correlated with its paired GT ROI with a correlation coefficient of more than 0.7, it was counted as a true positive *TP*. If it did not meet these conditions, it was counted as a false positive *FP*. A GT ROI not paired with any *TP* ROI obtained with another method was counted as a false negative (*FN*).

Under the above definitions of *TP*, *FP*, and *FN*, the precision, recall, and success rate were calculated as performance measures. Precision was defined as the proportion of true positives among the total number of ROIs obtained with other method, i.e. *Precision* = *TP/*(*TP* + *FP*), while recall was defined as the proportion of the number of GT ROIs paired with *TP* ROIs among the total number of GT ROIs, i.e. *Recall* = *TP/*(*TP* + *FN*). The success rate was defined as the harmonic mean between *Precision* and *Recall*.

## 3. Results

Here, we confirmed that LCCD efficiently works on data of high-density overlapping cells in comparison with CNMF and Suite2P. To demonstrate the efficacy of the temporal division and integration approach in LCCD, we also compared the performance of the blob detector with that of LCCD on both artificial and real data.

### 3.1. Performance evaluation with synthesized data

To quantitatively evaluate the performance of LCCD and the other methods, we used three samples of synthesized data of high-density overlapping cells in 256 × 256 pixels. The left-most left panel of Fig. 2A is the temporally maximum luminance image of one of the three samples. The other panels of Fig. 2A show ROIs obtained by LCCD, BD, CNMF and Suite2P from the sample. The numbers of ROIs obtained by LCCD were almost same as those obtained by CNMF and Suite2P, but not by BD, for the three samples of synthesized data (Fig. 2B).

**Figure 2:**
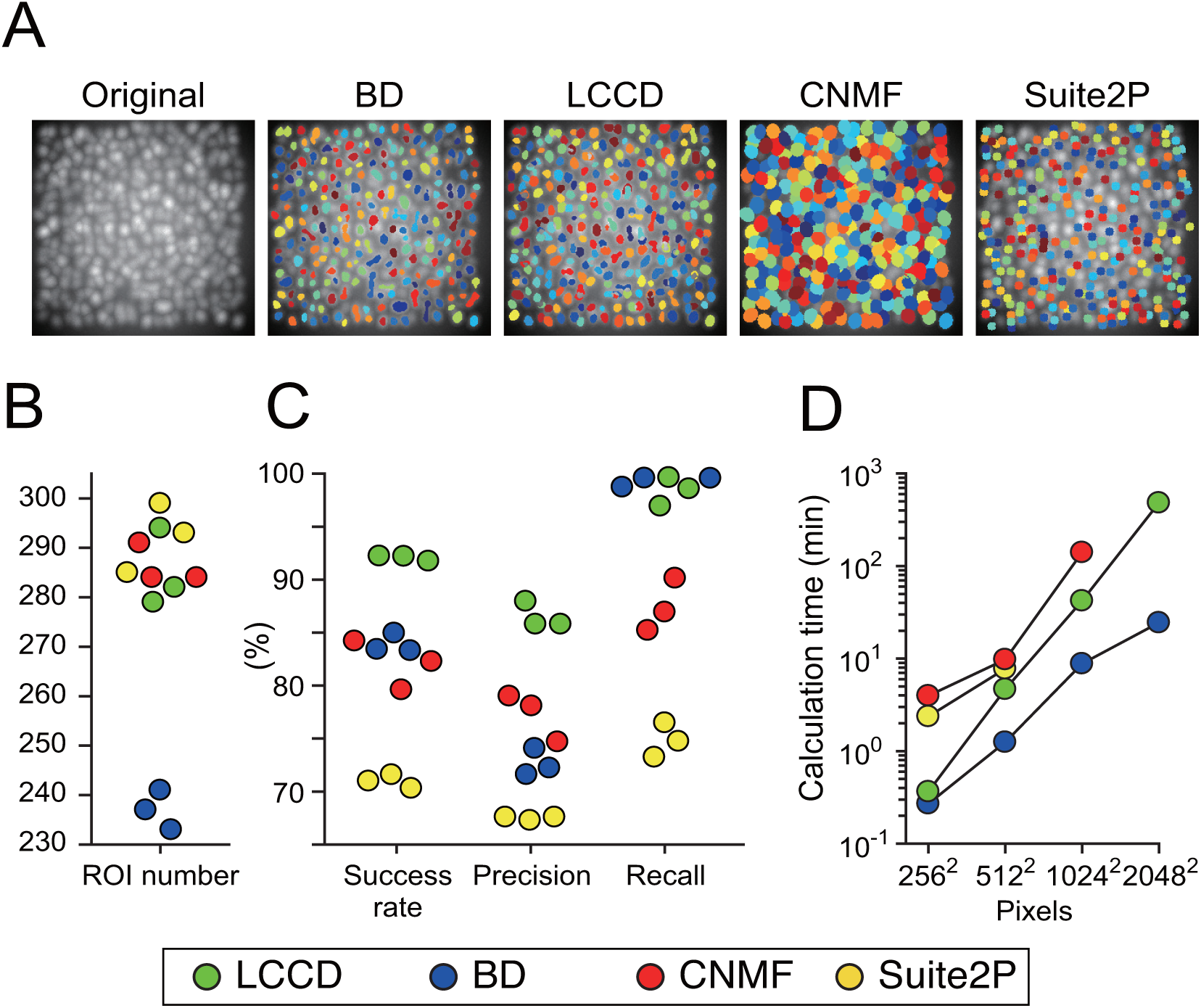
Comparison of BD, LCCD, CNMF and Suite2P using synthesized data of high-density overlapping cells in 256×256 pixels. (A) Temporal average of synthesized data and ROIs obtained with the four methods. (B) Total number of ROIs obtained with each method. The number of true ROIs is 324. (C) Precision, recall and success rate of each method. (D) Calculation time of each method as a function of the size of synthesized data: 225 cells in 256 × 256 pixels, 841 cells in 512 × 512 pixels, 3136 cells in 1024 × 1024 pixels, and 10000 cells in 2048 × 2048 pixels. The absence of a line extending to a pixel size case means that the process had not been completed even after three days on a high-performance computer.

To compare the performance of LCCD with those of BD, CNMF and Suite2P statistically with reference to the true ROIs, we calculated the precision, recall, and success rate. As shown in Fig. 2C, the success rate of Ex-LCCD was 92.1% on average, which was prominently the highest among the four methods. The success rates of BD and CNMF were almost equal, being respectively 83.9% and 82.0% on average and intermediate in value among the four methods. In contrast, the success rate of Suite2P was lowest among the four methods, and furthermore its precision and recall were lowest among the four methods.

Next, we evaluated the calculation time of the four methods processing four synthesized data: 225 cells in 256 × 256 pixels, 841 cells in 512 × 512 pixels, 3136 cells in 1024 × 1024 pixels, and 10000 cells in 2048 × 2048 pixels on the same high-performance computer (Fig. 2D). As shown in the figure, Suite2P did not complete its process on 1024 × 1024 or 2048 × 2048 pixels of data within three days, and the CNMF did not complete its process on 2048 × 2048 pixels of data within three days. On the other hand, LCCD and BD completed their processes on all four datasets. The calculation time of BD was over ten times shorter than those of CNMF and Suite2P, whereas the calculation time of LCCD was shorter than CNMF and Suite2P but longer than BD. For instance, BD and LCCD respectively took 9 minutes and 30 minutes to process 1024 × 1024 pixels of data, whereas CNMF took 140 minutes. Furthermore, BD and LCCD respectively took 24 minutes and 300 minutes to process 2048 × 2048 pixels of data, whereas the other methods had not completed the cell detection process within three days. Note that CNMF and Suite2P include heuristic pre- and post-processes that were not explained in much detail in their papers. CNMF and Suite2P spent much time on the pre-post processes. Thus, it is impossible to estimate the whole computational complexity of CNMF and Suite2P.

### 3.2. Performance evaluation with in-vivo data

We evaluated the performance of the top-three methods in the evaluation, LCCD, BD and CNMF, on in vivo data. To make CNMF work efficiently, we used two-photon calcium imaging data with a conventional field-of-view (256 × 256 pixels, 2000 frames). 18 samples were used.

Figure 3A shows ROIs obtained with LCCD, BD and CNMF from one of the 18 samples. Areas surrounded by white rectangles and the closeups in Figs. 3A indicate examples in which BD merged a few different cells into a single large ROI but LCCD and CNMF divided the same region into individual cell ROIs and examples in which LCCD and CNMF detected cells that BD failed to catch. Note that the initial number of ROIs in the CNMF algorithm was set 10% larger than the number of ROIs obtained with LCCD. The numbers of ROIs obtained with LCCD were significantly larger than those of BD for all samples, but were almost same as those of CNMF for the above initial condition (Fig. 3B).

**Figure 3:**
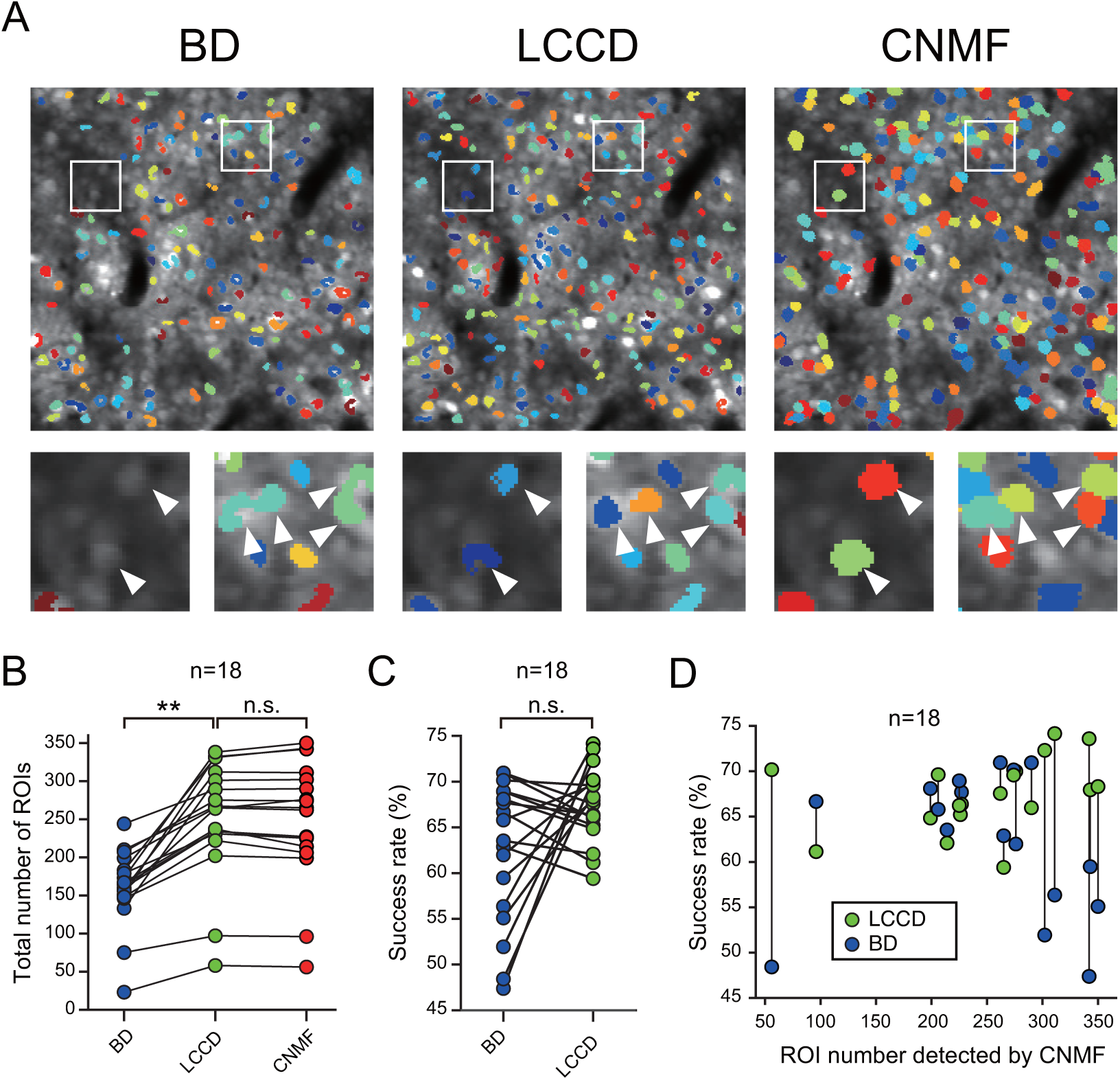
Performance evaluation of BD, LCCD and CNMF using in vivo data (256 × 256 pixels, 2000 frames) (A) ROIs obtained with the three methods. (B) Total number of ROIs obtained with each method for various samples. Each line in the graph indicates results of a method on a single sample. (C) Success rate of BD and LCCD with reference to CNMF for various samples. (D) Difference in success rate between BD and LCCD with reference to CNMF for various numbers of ROIs detected by CNMF.

Next, to quantitatively compare the performance of LCCD, BD and CNMF in situations where the true ROIs were not known, we measured the relative similarity of ROIs obtained with LCCD, BD, and CNMF by calculating the success rates of LCCD and BD with reference to CNMF. The success rates of LCCD and BD were respectively 67.5% and 62.5% on average. As shown in the profiles of the success rate depending on samples (Fig. 3C), there were samples for which the success rates of LCCD were almost the same or slightly lower than those of BD, whereas there were samples for which the success rates of LCCD were much larger than those of BD. The difference in success rate between LCCD and BD depended on the number of detected ROIs (Fig. 3D). There was a tendency whereby the success rate of LCCD became much larger than that of BD when the number of detected ROIs by CNMF was more than 300.

Next, we checked whether the LCCD ROIs that did not match those of CNMF were cells or not. Typical fluorescence time series of LCCD ROIs not matching those of CNMF showed transient activities (Fig. 4A), which are typical calcium kinetics of active cells (Chen et al., 2013). Using a spike detection algorithm based on the state-space model (Pnevmatikakis et al., 2016), we detected and counted transient events of each ROI that matched or did not match those of CNMF. Figure 4B shows the frequency distributions of the number of counted calcium transient events per ROI either matched or not matched to the ROIs of CNMF. 84.4% of non-matching ROIs had transient events, while 93.6% of the matching ROIs had transient events. The frequency of no matching ROIs without a transient event was 9% higher than that of matching ROIs. The profiles of the distributions were similar when the number of transient events was larger than 3.

**Figure 4:**
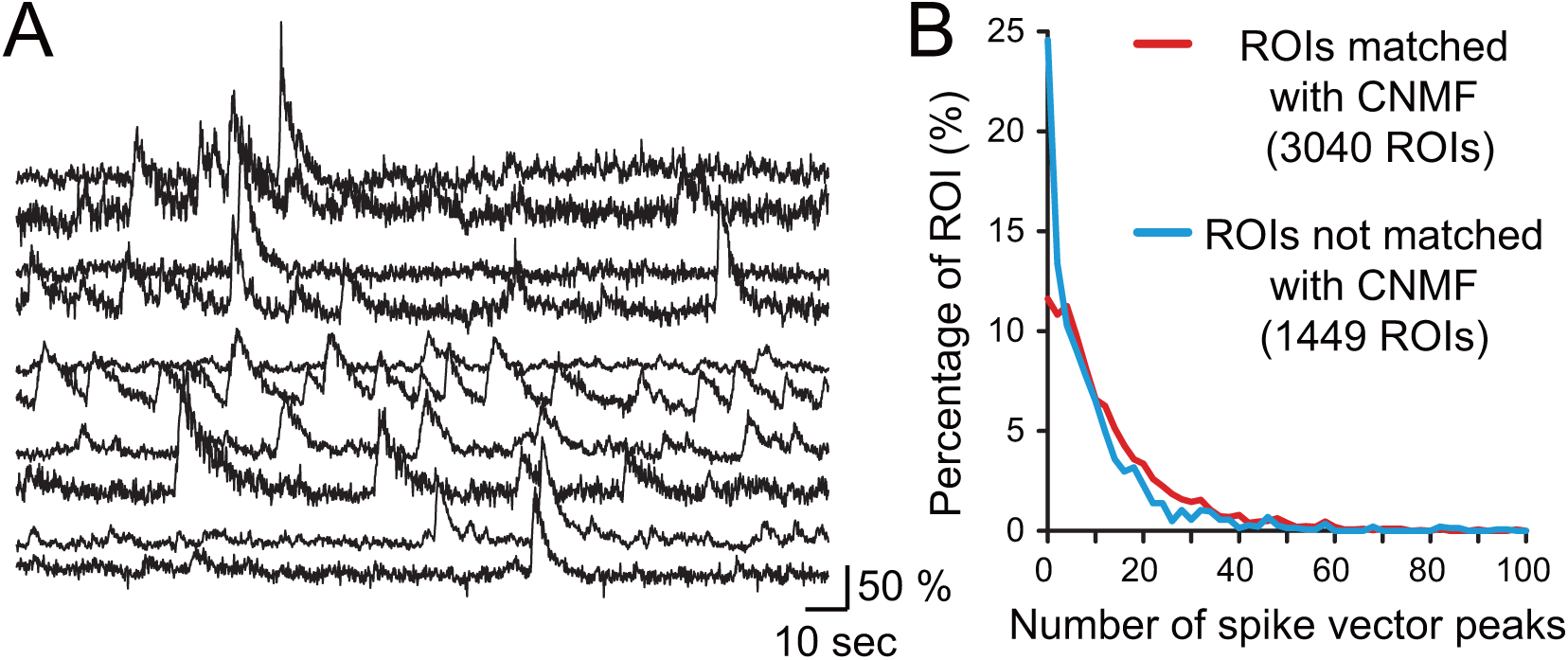
Profiles of fluorescence time series in LCCD ROIs not matched with those of CNMF. (A) Typical fluorescence time series of LCCD ROIs not matched with those of CNMF. (B) Distributions of the number of counted transient events per ROI matched or not matched with those of CNMF. TP: ROIs matched with those of CNMF. FP: ROIs not matched with those of CNMF.

## 4. Discussion

### 4.1. Summary of results and conclusion

The success rate of LCCD was prominently higher than BD and CNMF and Suite2P on the synthesized data (Fig. 2B). This result suggests that overlapping cells were separated into individual ones by using the temporal division and integration approach. Furthermore, we measured the relative similarity of the ROIs obtained with LCCD and CNMF on in-vivo data with a conventional FOV (256 × 256 pixels). The percentage of ROIs identified by LCCD that matched those of CNMF was on average 67.5%, as evaluated by calculating the success rate of LCCD relative to CNMF (Fig. 3C). In particular, the success rate of LCCD was higher than that of BD in the case of high-density GCaMP-expressing cells (more than 300 detected ROIs) (Fig. 3D). This result suggests that overlapping cells could be separated into individual ones as a result of using the temporal division and integration approach, even on in-vivo data. Moreover, 84.4% of the ROIs that did not match those of CNMF were confirmed to have calcium transient events (Fig. 4B). This result suggests that almost all of the LCCD ROIs that did not match those of CNMF were active cells. Furthermore, LCCD completed the cell detection task on 2048 × 2048 pixels of data within a reasonable amount of time (about 5 hours), whereas CNMF and Suite2P had not completed the cell detection process even after three days (Fig. 2D).

The conclusion obtained from these results is that LCCD makes it possible to detect cells from huge amounts of high-density imaging data within a practical time and with an accuracy comparable to or better than those of state-of-art methods such as CNMF and Suite2P.

### 4.2. Efficacy of temporal division and integration approach

Here, we point out two positive effects of the temporal division and integration approach on the performance of LCCD. As shown in Figs. 3A, there were cases in which BD merged a few different cells into a single large ROI, whereas LCCD divided up the same region into individual cell ROIs. If overlapping cells have different fluorescence variations over time, they have different active patterns, in which one is active and other one is inactive, in each of the short time frames. By integrating temporally different patterns of ROIs, overlapping cells can be separated into individual cell ROIs. As shown in Figs. 3A, there were cases in which LCCD detected cells that were not caught by BD. LCCD removes the background signal by using a long moving-average filter. However, the background signal intermittently changes with an amplitude comparable to variations in cell signals. Such an unsteady background cannot be removed by a long moving-average filter. On the other hand, the background can be more accurately removed within short time frames, because the background signal remains steady during a short period. Thus, the ability of cell detection is improved by the temporal division and integration approach.

### 4.3. Problem of LCCD and improvement method

The calculation time of LCCD was intermediate between that of BD and the other methods (Fig. 2D). In LCCD, BD is used to identify cells as ROIs in each of the divided frames, and these ROIs in temporally different frames are then integrated. The greatest bottleneck in this procedure is the ROI integration. Here, we think there is a possibility to speed up LCCD by improving ROI integration algorithm and its code.

## 5. Acknowledgements

This study was supported by the Cooperative Study Program of the National Institute for Physiological Sciences. The authors are grateful for support from the Japan Society for Promotion of Science, KAKENHI JP16H01539, JP16H01269 and JP19H04203 to T.A. This research was partially supported by the program for Brain Mapping by Integrated Neurotechnologies for Disease Studies (Brain/MINDS) from the Japan Agency for Medical Research and Development, AMED under the Grant Number JP19dm0207001. This research was also supported by a CREST grant (Grant Number JPMJCR1861) from the Japan Science and Technology Agency.

## Algorithm 1 Main routine of LCCD

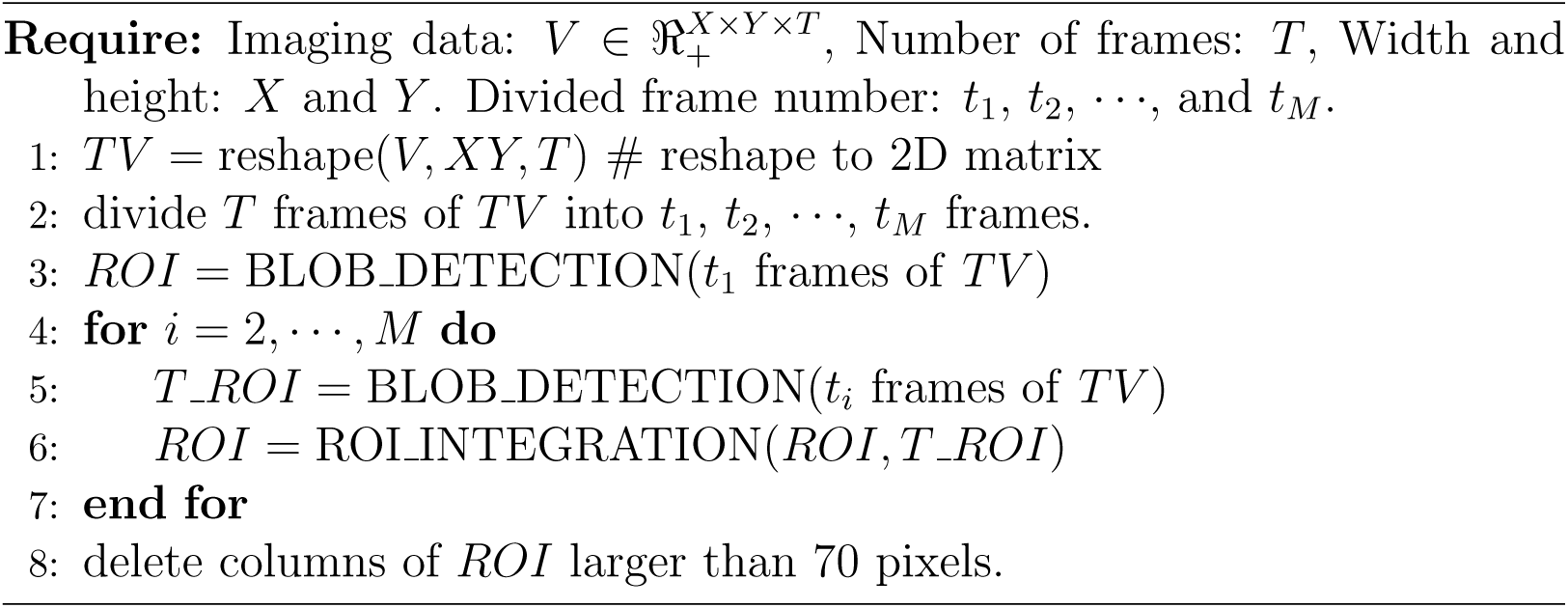

## Algorithm 2 blob detector (Matlab code is provided as a way to simplify the explanation)

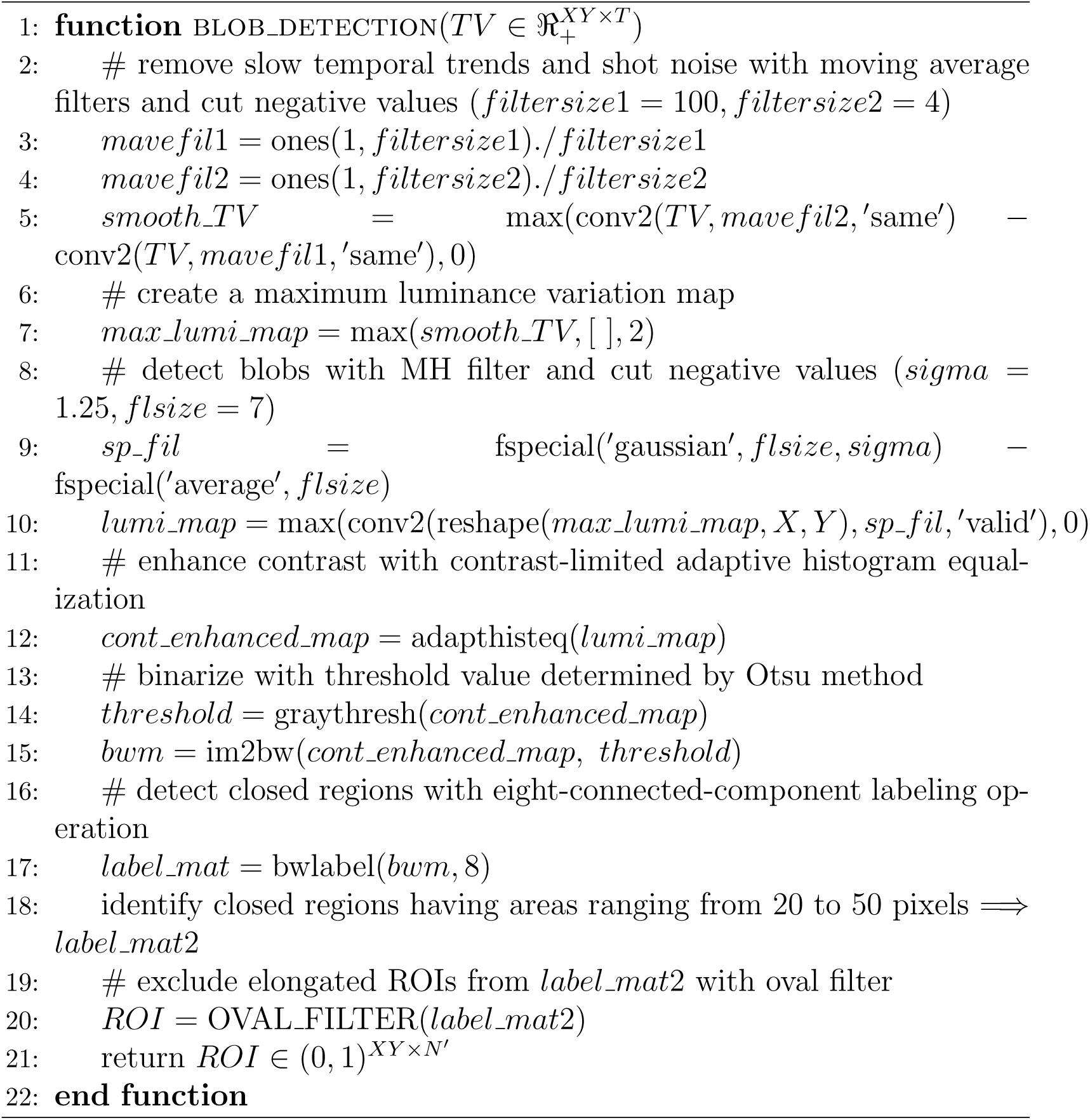

## Algorithm 3 Oval filter (Matlab code is provided as a way to simplify the explanation)

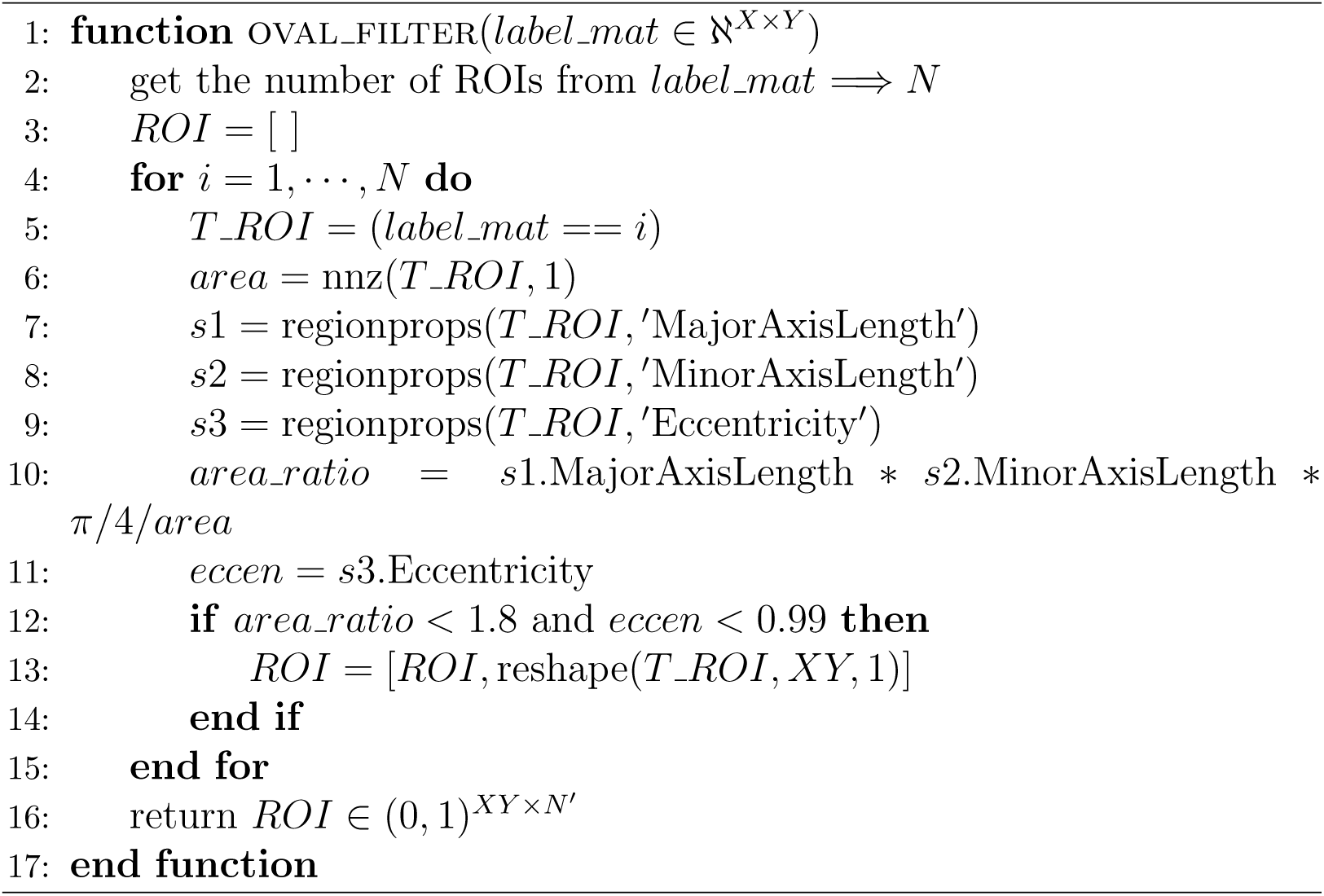

## Algorithm 4 ROI integration (Matlab code is provided as a way to simplify the explanation)

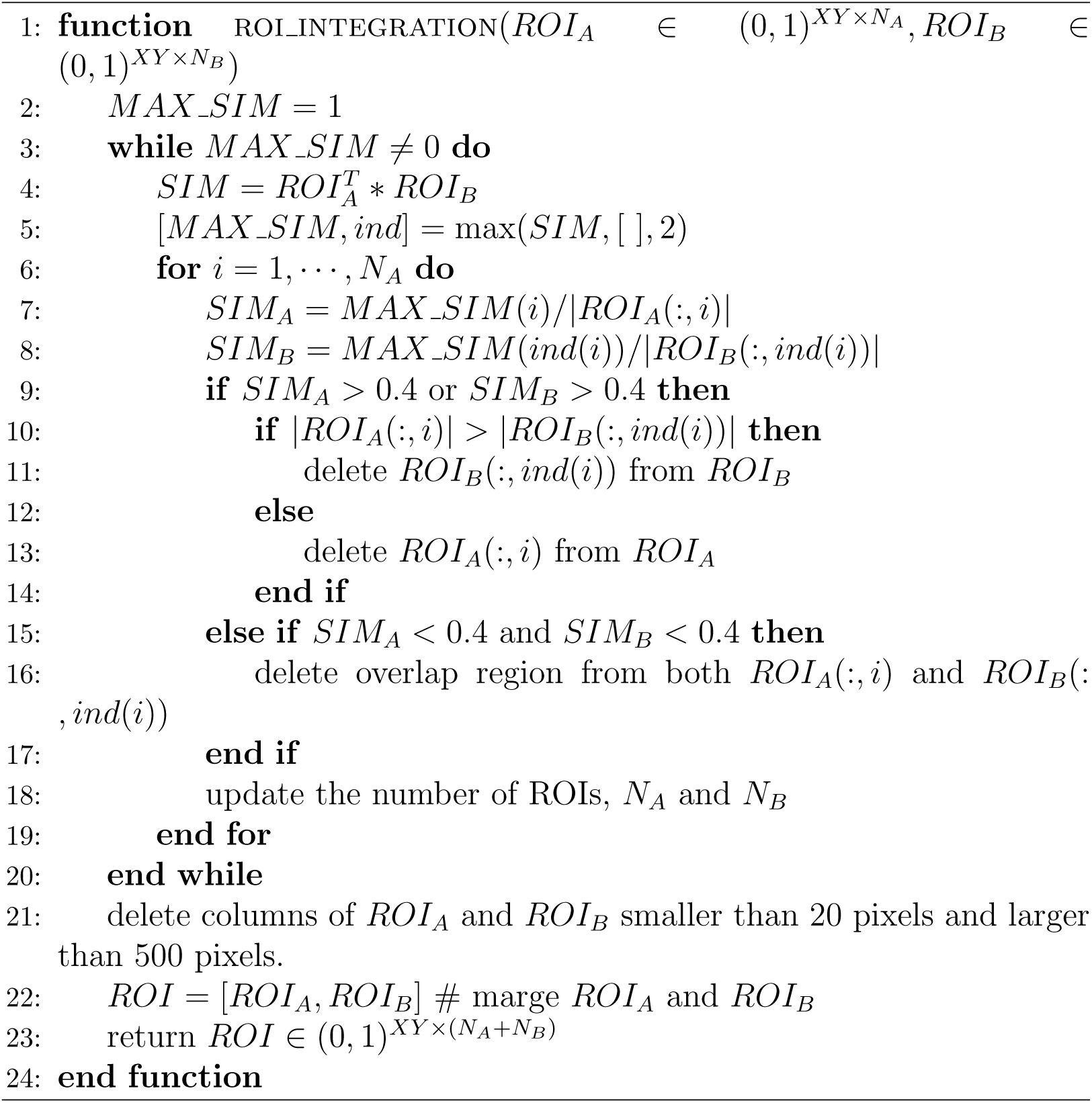

